# Third generation cephalosporin-induced L-form transition in *Shigella sonnei* reveals a virulence-survival trade-off underlying persistent infection

**DOI:** 10.1101/2025.07.17.665312

**Authors:** Sanjib Das, Arindam Mukherjee, Prolay Halder, Soumalya Banerjee, Supriya Mandal, Nivedita Roy, Ashis Debnath, Sreejani Banerjee, Manjistha Manna, Upama Mandal, Jiro Mitobe, Jeffery H. Withey, Asish Kumar Mukhopadhyay, Santasabuj Das, Debaki Ranjan Howlader, Hemanta Koley

## Abstract

Shigellosis remains a major global health burden, and the increasing prevalence of multidrug-resistant (MDR) *Shigella* strains is complicating effective antibiotic therapy. Bacteria may survive antibiotics by transitioning into cell wall-deficient L-forms, which are intrinsically resistant to β-lactams and can revert to a virulent state, potentially causing relapsing infections. Here we characterized a clinical MDR isolate *Shigella sonnei* HK8, a.k.a. PD552A, whose genome contains key resistance (*gyrA*, PBP3) and virulence (*icsA*) genes. Exposure to ceftriaxone induced a transition into a viable L-form state that was hyper-adhesive to macrophages *in vitro*. However, this survival adaptation was linked to a profound loss of pathogenicity. Using murine and guinea pig models, the L-form variant was shown to be profoundly attenuated, failing to cause the keratoconjunctivitis, diarrheal disease, or significant histopathology characteristic of the wild-type strain. These findings reveal a critical virulence-survival trade-off, positioning the L-form as a "stealth" phenotype that enables bacterial persistence at the expense of acute virulence. This offers a potential mechanism for asymptomatic carriage and recurrent infections, highlighting a previously underappreciated mechanism by which antibiotic treatment may resolve acute symptoms while permitting the persistence of a cryptic bacterial reservoir capable of driving recurrent infection.

**Importance:** Our research provides critical insight into the challenge of antibiotic treatment failure in shigellosis. By integrating experimental validation of reversible L-form transitions with mathematical modelling, we uncover a crucial virulence-survival trade-off. We show that MDR *S. sonnei* survives ceftriaxone by adopting a “stealth” phenotype, quantified by a high Stealth Index – maintaining bacterial burden while evading host inflammatory detection. These findings imply that standard antibiotics may resolve acute symptoms while inadvertently selecting for a cryptic, persistent reservoir poised for relapse. This work challenges the conventional view of therapeutic success and highlights an urgent need to develop novel diagnostic and therapeutic strategies capable of identifying and eliminating these resilient, "stealth" L-form persisters to achieve true bacterial clearance and prevent chronic infections.

## Introduction

Shigellosis, an intestinal infection caused by *Shigella* spp., is a leading bacterial cause of diarrheal morbidity and mortality worldwide, imposing a substantial burden on low- and middle-income countries (LMICs) [1, 2]. In these regions, fecal-oral transmission is facilitated by inadequate sanitation and a lack of potable water [3]. While high-income countries benefit from stringent regulatory oversight of food and water, they are not immune to the threat of drug-resistant *Shigella*. A 2023 CDC report revealed a significant increase in extensively drug-resistant (XDR) *Shigella* in the United States; a concerning trend linked to the overuse of broad-spectrum antimicrobials [5]. Cephalosporins, a class of β-lactam antibiotics, are frequently prescribed for shigellosis but can inadvertently select for resistant strains [6]. These agents inhibit peptidoglycan synthesis by binding to penicillin-binding proteins (PBPs), disrupting cell wall integrity and typically leading to cell death. However, under sublethal antibiotic pressure, bacteria can survive by transitioning into cell wall-deficient variants known as L-forms. This phenotypic switch renders them intrinsically resistant to cell wall-active antibiotics and alters their surface antigen profiles, allowing for evasion of host immune detection.

The capacity for such adaptation may underlie the phenomenon of chronic or persistent shigellosis. Although often presenting as an acute infection, studies have documented long-term carriage and transmission, particularly of *S. sonnei* in specific populations [10]. This persistence has been linked to plasmid-encoded structures, such as the O-antigen, that are retained even after antibiotic exposure, facilitating chronic carriage and posing a public health challenge [10]. The formation of L-forms represents a plausible mechanism for this persistence. L-forms of other Gram-negative bacteria evade the immune system by modifying pathogen-associated molecular patterns (PAMPs), such as lipopolysaccharide (LPS), thereby impairing recognition by host pattern recognition receptors (PRRs) and phagocytosis. Crucially, L-forms can revert to their virulent, cell-walled state once the antibiotic pressure is removed, potentially causing infection relapse. Despite the clinical relevance, the role of L-forms in *Shigella* pathogenesis remains poorly understood, partly due to the lack of robust animal models that fully recapitulate human shigellosis [9]. Therefore, it is imperative to investigate L-form biology in relevant experimental models to elucidate their contribution to antibiotic treatment failure and chronic infection.

Keeping these in mind, here we performed a comprehensive characterization of the clinical *Shigella sonnei* isolate HK8, a strain confirmed to be resistant to ceftriaxone and genotypically armed with a robust profile of antimicrobial resistance genes, toxins, and virulence factors. We demonstrated that exposure to this third-generation cephalosporin, while bactericidal, induces a survival strategy wherein the bacteria transform into a viable, cell wall-deficient L-form state. These L-forms exhibited an altered *in vitro* phenotype, characterized by heightened surface adhesion to RAW 264.7 macrophage cells. However, this adaptive change was accompanied by a profound loss of pathogenicity *in vivo*. In both systemic mouse infection and localized guinea pig models, the L-form variant was significantly attenuated, failing to induce the severe diarrheal pathology or keratoconjunctivitis characteristic of the wild-type strain. These findings highlight a critical virulence-survival trade-off in *Shigella*, opening new avenues for understanding bacterial persistence and the role of L-forms in chronic, sub-clinical infections.

## Results

To systematically investigate the relationship between antibiotic exposure, phenotypic switching, and virulence, we performed a combined genomic, phenotypic, and *in vivo* analysis of the clinical isolate *S. sonnei* HK8.

### Genomic Features and Phylogenetic Placement of *Shigella sonnei* HK8

Whole-genome sequencing of the clinical isolate *Shigella sonnei* HK8 revealed a draft genome of 4.57 Mb with a GC content of 50.56%, assembled into 105 contigs and predicted to contain 83 tRNAs and 4 rRNAs (Supplemental Figure S1A). Comprehensive whole-genome phylogenomic analysis confirmed the isolate’s taxonomic identity, demonstrating that HK8 forms a monophyletic clade with the *Shigella sonnei* ATCC 29930 type strain (Supplemental Figure S1B). This analysis also reaffirmed the well-established taxonomic complexity where *Shigella* species are phylogenetically nested within the broader *Escherichia coli* species complex. Consistent with its role as an enteric pathogen, *in silico* screening of the HK8 genome identified a large repertoire of putative virulence factors, distributed across categories of known and potential secreted and non-secreted effectors essential for pathogenesis (Figure 1A).

**Figure 1.**
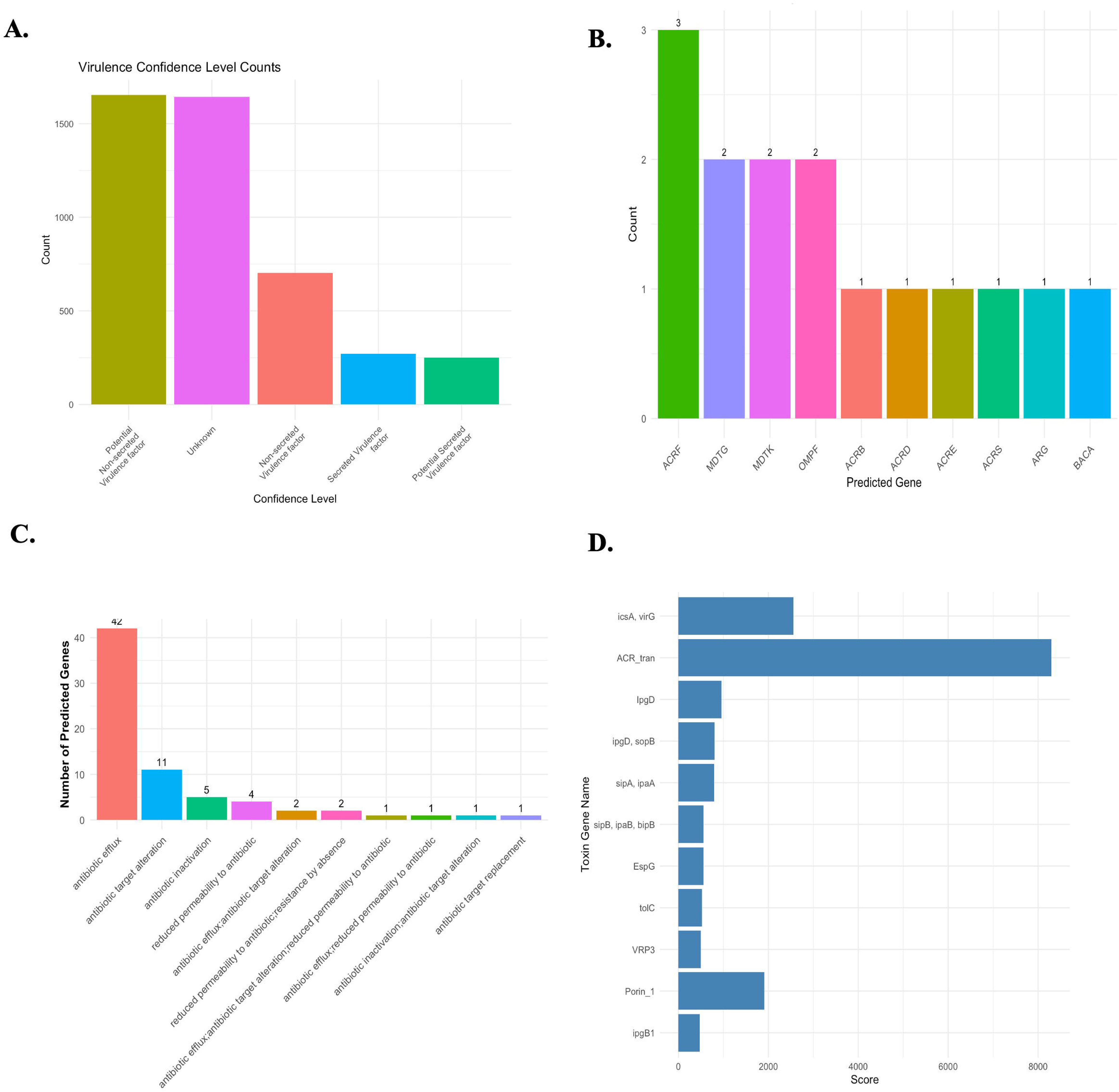
In Silico Profiling of Antimicrobial Resistance and Virulence Genes in *Shigella sonnei* HK8. **(A)** Bar chart showing the absolute counts of putative virulence-associated genes in the HK8 genome, categorized by confidence level. The analysis identified a large number of genes with homology to "Potential Non-secreted Virulence factors" and a substantial fraction classified as "Unknown." Genes with stronger evidence were categorized as "Non-secreted Virulence factors," "Secreted Virulence factors," and "Potential Secreted Virulence factors," providing a quantitative overview of the isolate’s pathogenic potential. **(B)** A bar chart showing the frequency of the top 10 most abundant predicted AMR genes in the HK8 genome. The analysis indicates a high prevalence of genes encoding components of multidrug efflux pumps, such as *ACRF* (3 copies), as well as genes *MDTG, MDTK,* and *OMPF* (2 copies each). **(C)** A bar chart categorizing the predicted AMR genes based on their mechanism of action. Antibiotic efflux is the most dominant mechanism, with 42 distinct genes identified. Other notable mechanisms include antibiotic target alteration (11 genes) and antibiotic inactivation (5 genes). **(D)** A horizontal bar chart ranking the top 20 predicted toxin and virulence-associated genes based on their prediction score from computational analysis. The highest-scoring hits include ACR_tran (part of the AcrAB-TolC efflux system, linked to both resistance and virulence) and the *icsA, virG* locus, essential for intracellular actin-based motility. Other key *Shigella* virulence factors such as *ipgD, sipA/ipaA*, and *ipgB1* are also highly ranked, underscoring the pathogenic potential of the isolate.

### Multidrug-Resistant Phenotype and Its Genetic Determinants

Phenotypic characterization via antibiotic disc diffusion assays confirmed that *S. sonnei* HK8 possesses a clinically significant multidrug-resistant (MDR) profile. The isolate exhibited high-level resistance to multiple antibiotic classes, including β-lactams (ampicillin, ceftriaxone), aminoglycosides (kanamycin, neomycin), fluoroquinolones (ciprofloxacin), and chloramphenicol (Supplementary Table 1). To elucidate the genetic underpinnings of this phenotype, the genome was interrogated for known antimicrobial resistance genes (ARGs) and resistance-conferring mutations. This genomic analysis identified several critical single nucleotide polymorphisms (SNPs) in key resistance-associated genes (Supplementary Table 2). Specifically, a D87G substitution was found in *gyrA*, a classic mutation known to confer resistance to fluoroquinolones. Furthermore, two mutations were identified in *PBP3* (D350N, S357N), the transpeptidase enzyme and primary target of β-lactam antibiotics; these alterations likely reduce the binding affinity of ceftriaxone, explaining the observed resistance. Additionally, mutations in the global transcriptional regulator *marR* were identified, which often leads to the overexpression of multidrug efflux pumps and contributes to the broad resistance profile observed in this clinical isolate.

### In Silico Analysis Reveals a Genetic Basis for Multidrug Resistance and Virulence

To thoroughly elucidate the genetic underpinnings of the observed multidrug-resistant phenotype, we systematically interrogated the annotated *S. sonnei* HK8 genome for known antimicrobial resistance (AMR) and virulence-associated genes (Figure 1B – D). The analysis of AMR determinants confirmed that antibiotic efflux is the principal resistance strategy encoded in the genome, with 42 distinct genes assigned to this category (Figure 1C). Efflux pumps are known to contribute significantly to intrinsic and acquired multidrug resistance. This is strongly supported by the high frequency of genes encoding efflux pump components, with ACRF (3 copies) being most prevalent, followed by MDTG, MDTK, and the outer membrane porin OMPF (2 copies each) (Figure 1B). Other significant resistance strategies, such as antibiotic target alteration (11 genes) and enzymatic antibiotic inactivation (5 genes), were also identified. This diverse genetic arsenal provides a clear molecular basis for the broad-spectrum resistance profile observed phenotypically.

In addition to resistance determinants, the genome of HK8 encodes a comprehensive suite of virulence factors essential for orchestrating shigellosis (Figure 1D). Critically, the analysis identified the canonical virulence factor *icsA* (virG), which is indispensable for the intracellular actin-based motility required for cell-to-cell spread. Furthermore, key components and effectors of the Type III Secretion System (T3SS), a molecular syringe used to inject proteins into host cells, were highly ranked, including *ipgD, sipA/ipaA,* and *sipB/ipaB.* These effectors are crucial for mediating host cell invasion and manipulating cellular processes. Interestingly, the efflux pump component ACR_tran received the highest prediction score, highlighting a critical dual function for this system in both antibiotic resistance and pathogenesis, likely by expelling host-derived antimicrobial compounds.

### Ceftriaxone Treatment Inhibits Growth and Induces a Filamentous Phenotype in *S. sonnei* which converts back into wild type when the drug is taken away

To characterize the direct physiological and morphological responses of *S. sonnei* HK8 to β-lactam pressure, we first evaluated its growth kinetics in the presence of a sub-inhibitory concentration of ceftriaxone (5 µg/mL). The resulting growth analysis demonstrated that ceftriaxone exposure significantly impaired bacterial proliferation compared to the robust growth of the untreated control culture (Figure 2A). This bacteriostatic effect was characterized by a notably extended lag phase, suggesting a period of cellular adaptation to the antibiotic stress, followed by a much slower rate of growth that resulted in a substantially lower final optical density (OD600) after 8 hours (p < 0.001).

**Figure 2.**
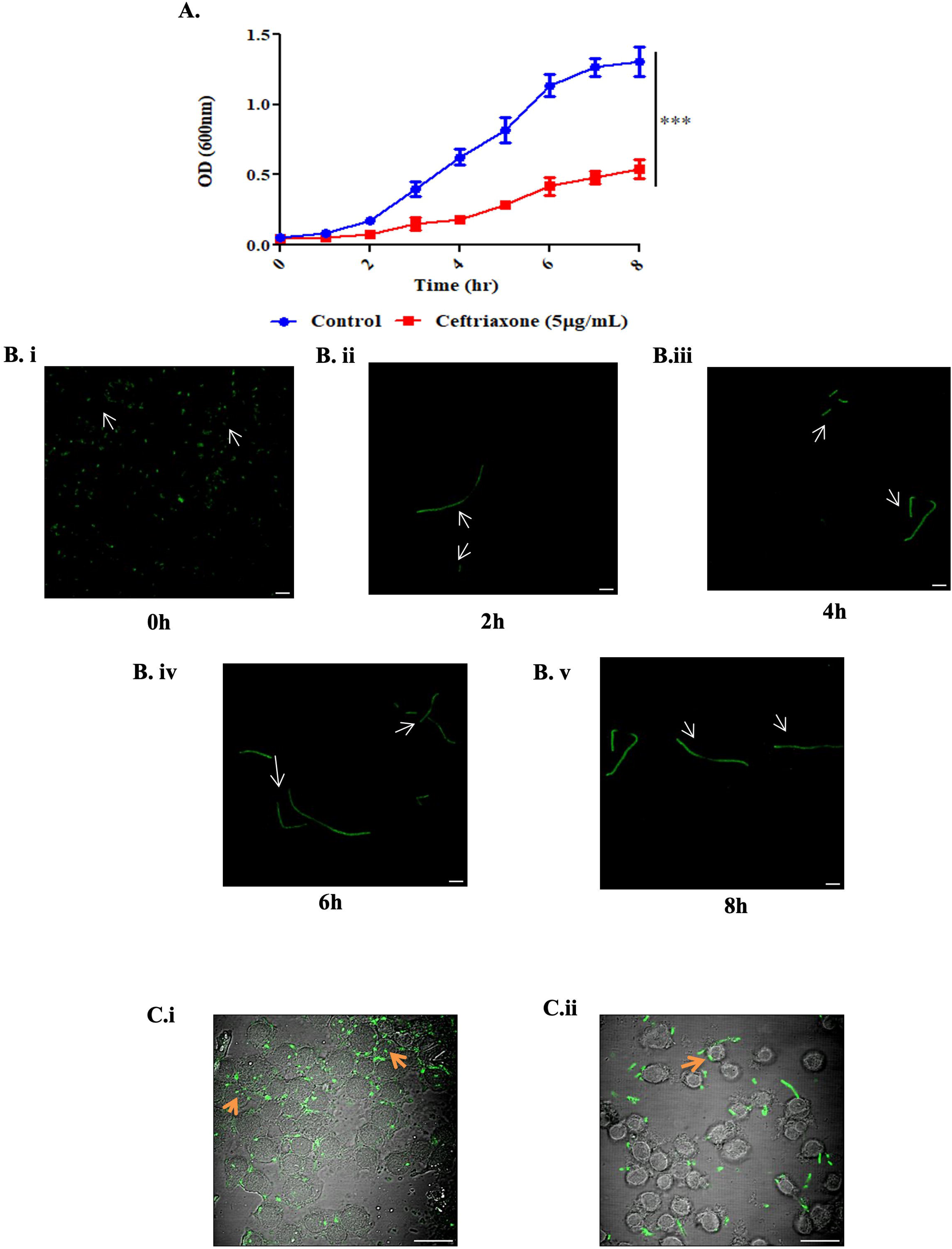
Ceftriaxone-Induced L-form Transformation of *S. sonnei* and Altered Interaction with Murine Macrophages. **(A)** Quantification of viable *S. sonnei* following a 24-hour exposure to a gradient concentration of ceftriaxone (∼MIC). The treatment resulted in a significant multi-log reduction in culturable bacteria (CFU/mL) compared to the untreated control yet allowed for the persistence of a resistant subpopulation. Data are presented as mean ± SD from three replicates. \*\*\**p* < 0.001. **(B & C)** Confocal microscopy images visualizing the interaction between *S. sonnei* and the RAW 264.7 macrophage cell line. **(B)** In the untreated control condition, classical, walled bacteria (green) are observed primarily populating the junctions between macrophage cells. **(C)** Following treatment with ceftriaxone, the surviving bacteria undergo a morphological shift into large, cylindrical L-forms (green). These L-forms exhibit a markedly different interaction dynamic, showing heightened and close association with the macrophage cell body. This observation supports the role of macrophages in facilitating the L-form state and suggests a shift in host-pathogen interaction that may promote persistence. Bar in figure 2.B: 5 μm, 2.C: 20 μm.

Concurrent with this growth inhibition, we observed profound morphological alterations using fluorescence microscopy to track bacterial cell shape over time (Figure 2B). While bacteria in the control culture consistently maintained their typical coccobacillary shape, those exposed to ceftriaxone failed to complete cell division and instead began to form long, multinucleated filaments. This filamentous phenotype, a classic hallmark of β-lactam-induced stress resulting from the inhibition of PBP-mediated peptidoglycan synthesis required for septation, became progressively more pronounced over the 8-hour period. Significant cell elongation was visible as early as 2 hours post-exposure, culminating in extensive, filaments by 8 hours (Figure 2B, ii-v).

Given this dramatic change in cell morphology, we next sought to determine if it affected the bacteria’s interaction with host immune cells by co-culturing them with the RAW 264.7 macrophage cell line (Figure 2C). In the control condition, the classical, rod-shaped bacteria displayed limited direct contact with the macrophage cell surface, tending instead to populate the junctions between cells (Figure 2C.i). In stark contrast, the ceftriaxone-induced filamentous bacteria exhibited a completely altered interaction dynamic. These elongated cells showed a strong propensity to adhere directly and extensively across the surfaces of the macrophage cell bodies (Figure 2C.ii). This suggests a significant shift in the pathogen’s surface properties and host-pathogen interaction, where sublethal antibiotic exposure promotes a hyper-adhesive phenotype. This may represent an alternative persistence strategy, facilitating surface colonization when classical T3SS-mediated invasion mechanisms are compromised.

To definitively validate the L-form state and the observed reversion, we utilized a SYTO 9/Propidium Iodide (PI) dual-staining assay (Figure 3). Wild-type *S. sonnei* cells successfully excluded PI due to an intact cell envelope, appearing uniformly green (Figure 3A). In contrast, the ceftriaxone-induced filamentous variants demonstrated significant PI uptake (Figure 3B), providing functional evidence of a compromised or absent peptidoglycan barrier. Crucially, upon removal of the antibiotic, these variants reverted to a vegetative morphology and completely restored their PI-exclusion properties within one hour (Figure 3C). This rapid restoration of the structural barrier distinguishes these viable, persistent L-forms from the irreversibly damaged, PI-saturated cells in the ethanol-treated control group (Figure 3D), confirming that the ceftriaxone-induced transition is a reversible survival strategy.

**Figure 3.**
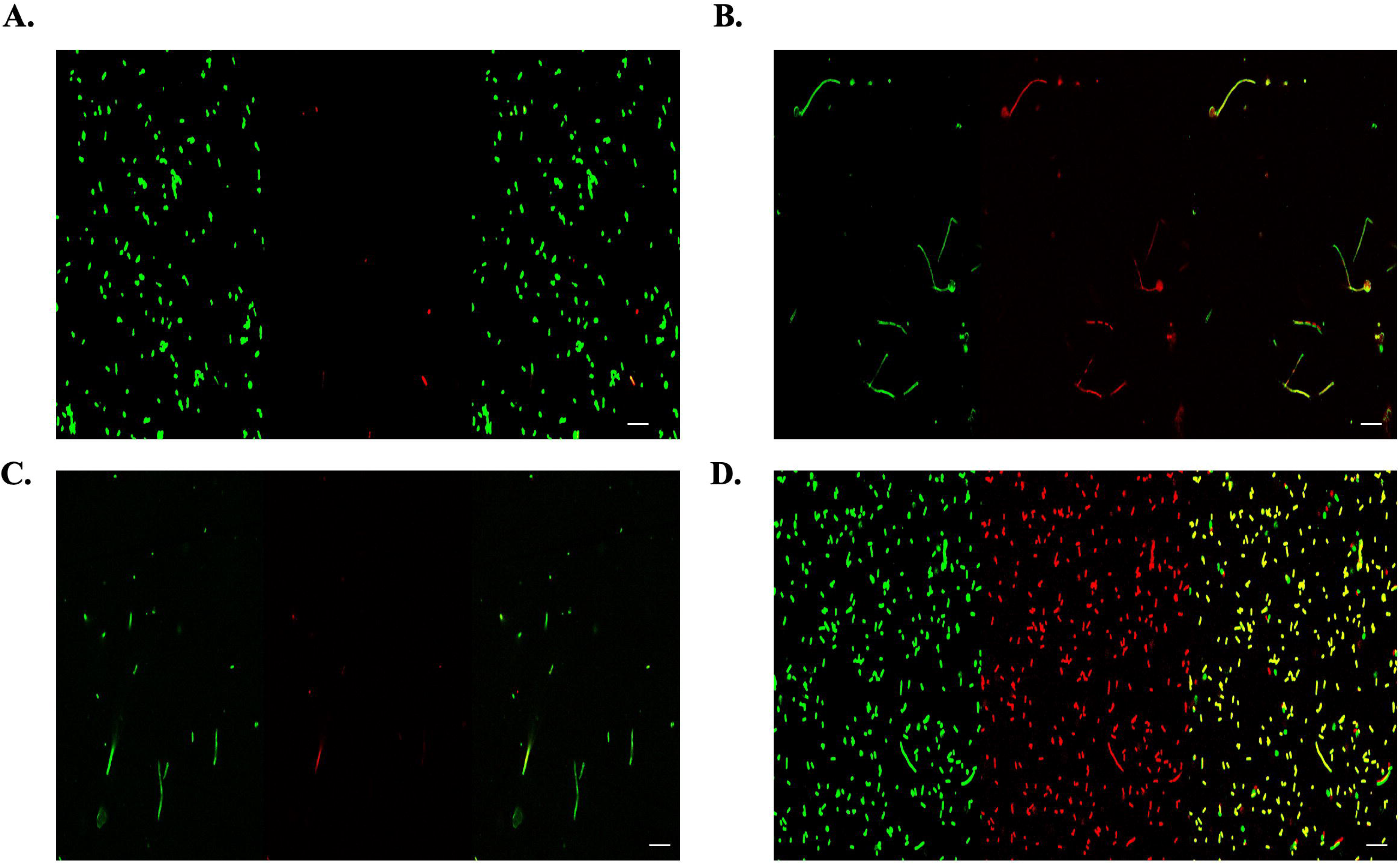
Reversible Membrane Permeability and Viability Assessment of *S. sonnei* L-forms. Representative confocal microscopy images using a SYTO 9/Propidium Iodide (PI) dual-staining assay to evaluate cell envelope integrity across different phenotypic states. Images were processed using Fiji (ImageJ); raw hyperstacks were separated into channels, converted using the ‘Stack to RGB’ function, and arranged into three-panel montages. Within each lettered panel (A–D), the side-by-side segments represent the same field of view showing the green channel (SYTO 9; all cells), the red channel (PI; membrane-compromised cells), and the merged image (yellow), respectively. **(A)** Wild-type *S. sonnei* displaying a typical rod-shaped vegetative morphology and successful exclusion of PI, indicating an intact cell envelope. **(B)** Ceftriaxone-induced L-forms exhibiting significant filamentous elongation and PI uptake, providing functional evidence of a compromised or absent peptidoglycan barrier in the L-form state. **(C)** Revertant bacteria one hour after the removal of ceftriaxone; the cells have rapidly restored their vegetative morphology and PI-exclusion properties, demonstrating the restoration of a functional structural barrier. **(D)** Ethanol-treated (70%) control group showing saturated PI uptake and irreversible membrane damage, distinguishing dead cells from the viable, persistent L-forms shown in panel B. Scale bars represent 5 μm.

### L-form *S. sonnei* Exhibits Attenuated Virulence in Animal Models of Infection

To determine if the phenotypic transition to the L-form state impacts the bacterium’s pathogenic potential *in vivo*, we compared its virulence against the wild-type (WT) strain using two complementary animal models of shigellosis (Figure 4).

**Figure 4.**
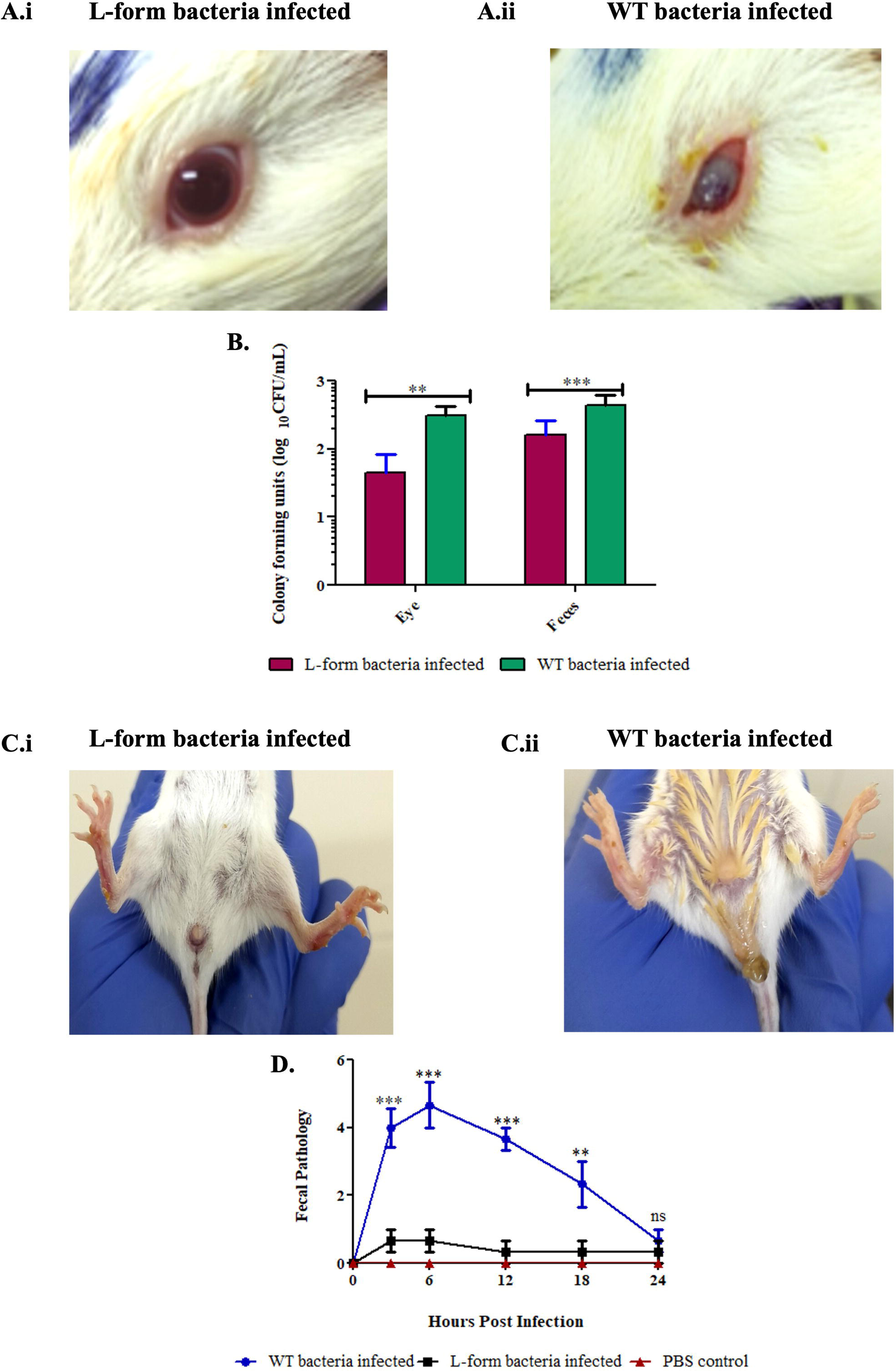
Attenuated Virulence of *Shigella sonnei* L-forms in *In Vivo* Models of Shigellosis. **(A)** Representative images from the Sereny test in guinea pigs 48 hours post-infection. **(A.i)** The eye infected with L-form bacteria demonstrates no significant signs of inflammation or pathology. **(A.ii)** In contrast, the eye infected with WT bacteria displays severe keratoconjunctivitis, characterized by inflammation and mucopurulent discharge, confirming the virulence of the classical form. **(B)** Quantification of bacterial burden recovered from the eyes (48 hours post-infection) and feces (6 hours post-infection). In both sites, the number of colony-forming units (CFU) recovered from L-form-infected animals was significantly lower than from those infected with the WT strain (**p < 0.01 for eyes, ***p < 0.001 for feces). **(C)** Representative images of BALB/c mice 6 hours after intraperitoneal infection. **(C.i)** The mouse infected with L-form bacteria appears healthy, with no visible signs of distress or diarrhea. **(C.ii)** The mouse infected with WT bacteria exhibits clear signs of diarrhea, indicative of acute gastrointestinal pathology. **(D)** Fecal pathology scores monitored over 24 hours post-infection. Infection with WT bacteria induced a rapid and significant peak in pathology at 6 hours, which gradually resolved. In contrast, L-form infection induced no significant pathological symptoms compared to the PBS control group.

First, we employed the guinea pig Sereny test, a classic model for assessing the ability of *Shigella* to invade corneal epithelial cells and elicit a localized inflammatory response. Infection with the WT strain elicited a robust, acute keratoconjunctivitis within 48 hours, characterized by severe conjunctival inflammation, opacity, and mucopurulent discharge (Figure 4A.ii). In stark contrast, the conjunctiva of animals infected with the L-form variant remained clear and quiescent, showing no overt signs of pathology (Figure 4A.i). This striking lack of clinical disease directly correlated with a diminished capacity to establish a productive infection, as significantly fewer colony-forming units (CFUs) were recovered from the eyes of L-form infected animals compared to those infected with the WT strain (p < 0.01) (Figure 4B).

Next, we assessed systemic disease progression and gastrointestinal involvement using an intraperitoneal (I.P.) infection model in BALB/c mice. Infection with wild-type *S. sonnei* induced a rapid onset of fulminant disease, manifesting as acute diarrheal symptoms, a key clinical indicator in this model (Figure 4C.ii). The severity of this condition was quantitatively captured by a fecal pathology score, which peaked sharply and significantly at 6 hours post-infection for the WT group before gradually resolving (p < 0.001) (Figure 4D). In contrast, mice challenged with the L-form variant remained completely asymptomatic throughout the 24-hour observation period, exhibiting no clinical signs of diarrhea or distress (Figure 4C.i). Accordingly, their fecal pathology scores were statistically indistinguishable from the PBS-injected negative control group. This clinical quiescence was mirrored by the bacteriological findings, as the bacterial load recovered from the feces of L-form infected mice was significantly lower than in the WT group (p < 0.001) (Figure 4B).

Collectively, these *in vivo* experiments strongly demonstrate that while the wild-type *S. sonnei* strain is highly virulent, its L-form variant is profoundly attenuated, with a severely compromised ability to colonize host tissues, trigger inflammation, and cause disease in both localized and systemic contexts.

### L-form *S. sonnei* Fails to Induce Significant Enterotoxicity and Histopathology in the Guinea Pig Ileal Loop Model

To directly assess the enterotoxic potential and tissue-damaging capabilities of the different bacterial forms, we employed the guinea pig ligated ileal loop model, a gold standard for studying enteric pathogenesis. In this assay, loops were inoculated with wild-type (WT) *S. sonnei*, its L-form variant, *Vibrio cholerae* as a positive control for severe enterotoxicity, and PBS as a negative control.

After 18 hours, gross examination of the loops revealed pronounced differences (Figure 5A). As expected, the positive control loop inoculated with *V. cholerae* exhibited severe hemorrhagic fluid accumulation, characteristic of its potent enterotoxin. In contrast, loops containing WT *S. sonnei* and the L-form variant showed substantially less fluid accumulation.

**Figure 5.**
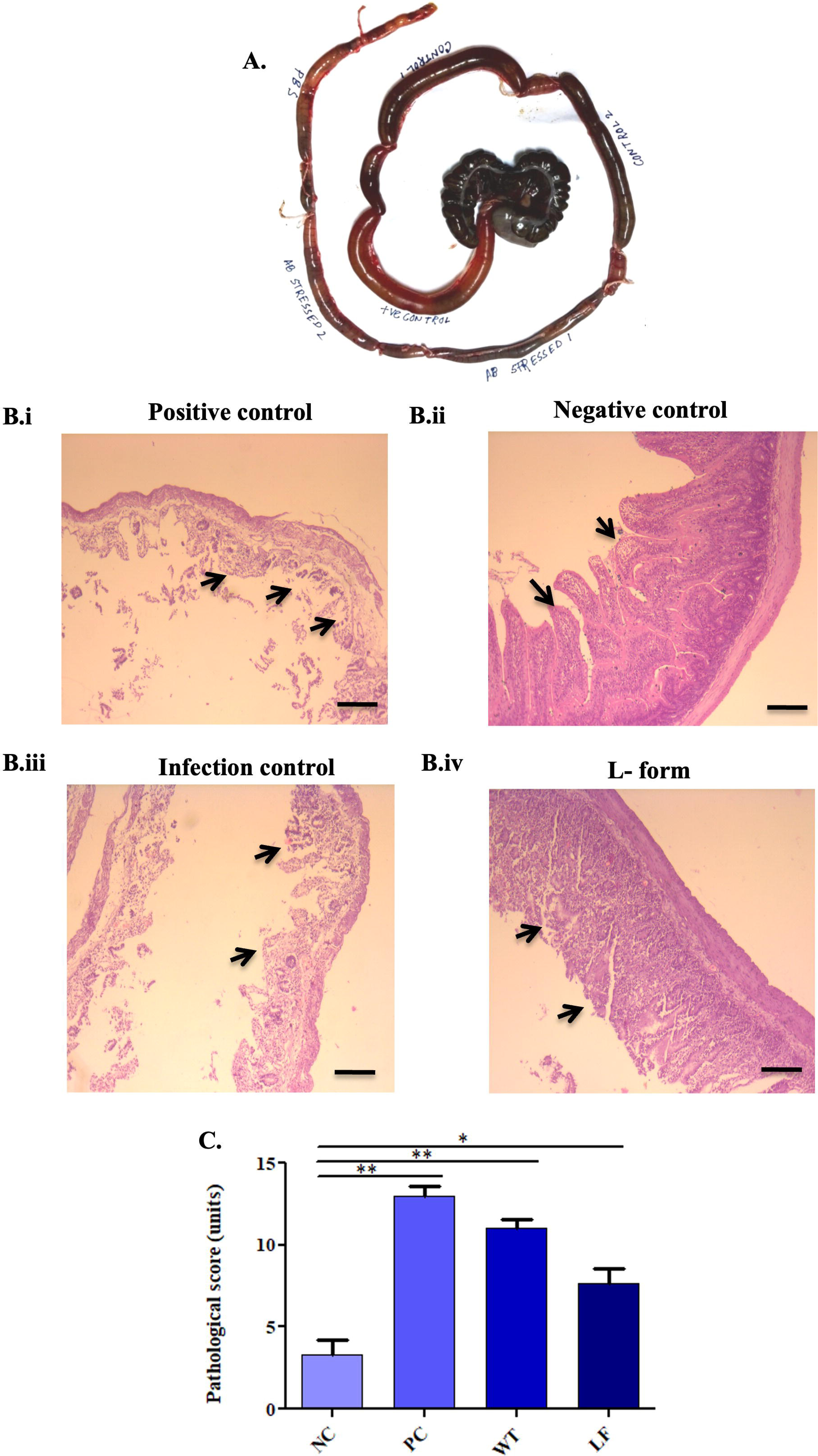
Attenuated Enterotoxicity and Histopathology of *Shigella sonnei* L-forms in the Guinea Pig Ileal Loop Model. Assessment of *S. sonnei* enterotoxicity across wild-type (WT) and L-form phenotypes via measurement of fluid accumulation and histopathology in the guinea pig ileal loop model (18 h post-infection). **(A)** Gross morphology of representative ligated ileal loops. The central loop, inoculated with the positive control (*Vibrio cholerae*), exhibits severe fluid accumulation and hemorrhagic damage. In contrast, loops inoculated with other agents show varying degrees of pathology. **(B)** Representative hematoxylin and eosin (H&E) stained sections of the ileal mucosa from each experimental group. **(B.i)** Positive Control (*Vibrio cholerae*): Demonstrates severe histopathological damage, including extensive epithelial shedding, loss of villous structure, and inflammatory cell infiltration (arrows). **(B.ii)** Negative Control (PBS): Displays normal, intact histological architecture with well-preserved villi (arrows). **(B.iii)** WT *S. sonnei* Infection: Exhibits significant pathological changes characteristic of shigellosis, including blunted villi, loss of epithelial integrity, and inflammatory infiltrates. **(B.iv)** L-form Infection: The intestinal surface topology is largely preserved, with minimal epithelial damage and significantly reduced inflammation compared to the WT infection. **(C)** Quantitative histopathological scoring of the ileal loop tissues. The scores confirm that both the positive control (PC) and WT *S. sonnei* induced significantly higher pathology than the negative control (NC) and L-form (LF) groups. Notably, the pathological score for the L-form group was significantly lower than that of the WT group, indicating attenuated tissue-damaging potential. Data are presented as mean ± SD. Asterisks denote statistical significance (*p < 0.05, **p < 0.01).

Histopathological examination of H&E-stained tissue sections provided a microscopic view of the damage (Figure 5B). The negative control loop (PBS) displayed a healthy, normal intestinal architecture with intact villi (Figure 5B.ii). Conversely, the positive control (*V. cholerae*) showed extensive necrosis and complete destruction of the villous structure (Figure 5B.i). The loop infected with WT *S. sonnei* presented significant pathology characteristic of shigellosis, including blunting and fusion of villi, loss of epithelial barrier integrity, and substantial infiltration of inflammatory immune cells into the lamina propria (Figure 5B.iii). Remarkably, the loop inoculated with the L-form variant showed a dramatically different and much milder outcome. The intestinal surface topology was largely preserved, with minimal epithelial disruption and significantly reduced inflammatory infiltration, closely resembling the negative control tissue (Figure 5B.iv).

These qualitative observations were confirmed through a quantitative histopathological scoring system (Figure 5C). The analysis verified that both the positive control and WT *S. sonnei* groups induced significantly higher pathology scores compared to the negative control (p < 0.01). Critically, the pathological score for the L-form group was significantly lower than that of the WT group (p < 0.05), further supporting the conclusion that the L-form of *S. sonnei* is profoundly attenuated in its ability to cause the acute inflammatory tissue damage typical of shigellosis.

### Mathematical Simulation Reveals a Therapeutic Tipping Point for L-form Persistence

To integrate our *in vitro* and *in vivo* findings, we employed a compartmental mathematical model to simulate the dynamics of the virulence-survival trade-off. Our simulations, calibrated to observed growth rates (Fig 2A), demonstrated a clear "therapeutic tipping point" at sub-inhibitory ceftriaxone concentrations. At these levels, the model predicted a rapid collapse of the classical virulent population (B_c_) and a concurrent spike in the cell-wall-deficient reservoir (B_l_). Furthermore, by applying the host pathology function (P), we calculated a stealth index (Σ) for both bacterial states. While WT infections exhibited a low Σ due to high symptomatic damage, the transitioned L-form population achieved a significantly higher index. This suggests that the L-form transition is not merely a survival mechanism but an optimized stealth strategy that allows for a substantial bacterial load to persist in host tissues while remaining below the threshold of inflammatory detection observed in our murine and guinea pig models (Fig 4 and 5). These dynamics are captured in our integrative model, which highlights the time-resolved emergence of the stealth reservoir and the resulting spike in the Stealth Index (Fig. 6A-C).

**Figure 6.**
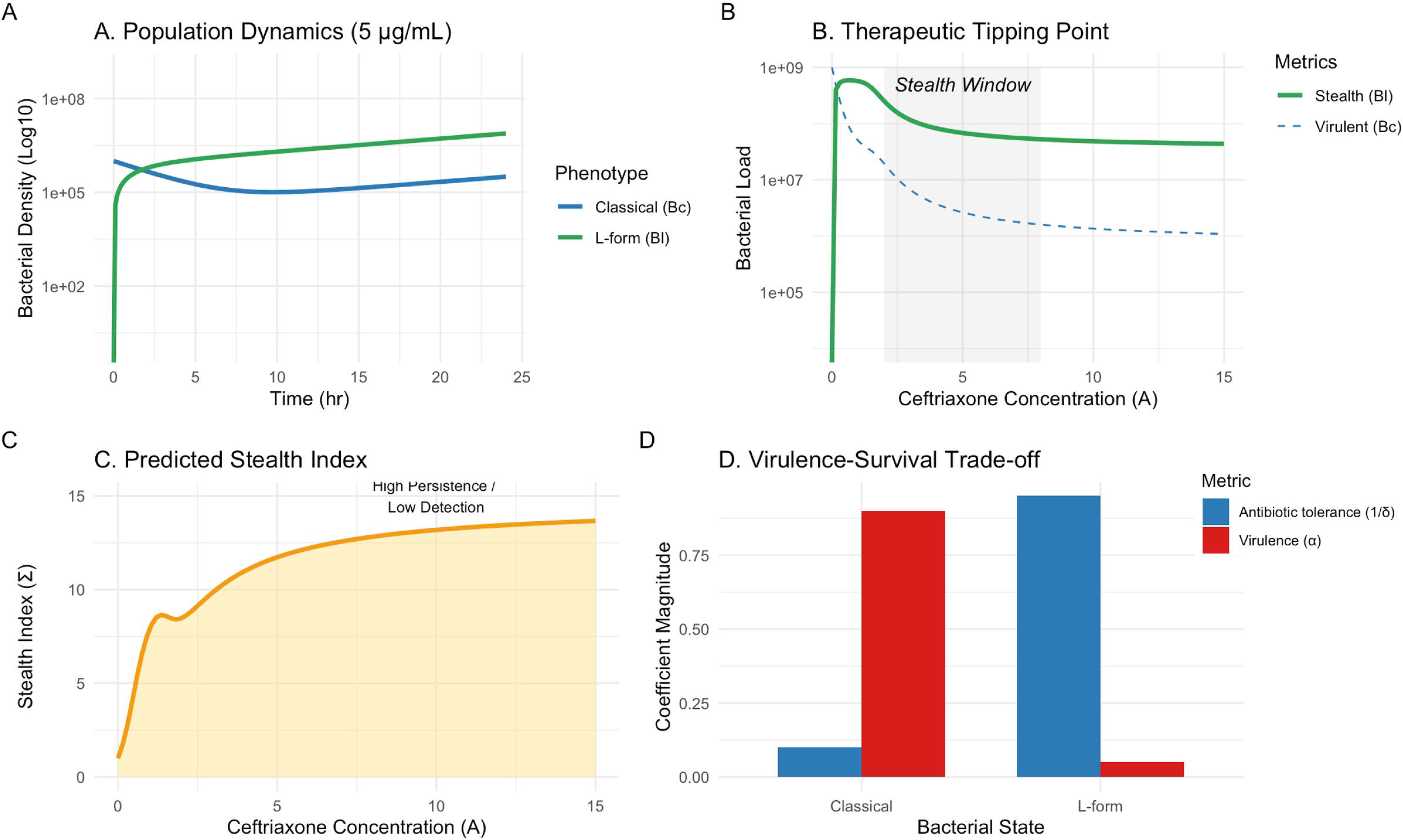
Simulation of antibiotic-driven phenotypic switching and stealth persistence in *Shigella sonnei*. **(A)** Time-course simulation of the transition from virulent *Bc* (blue) to persistent *Bl* (green) under antibiotic pressure. Population values represent normalized simulated bacterial densities derived from the ODE model rather than direct CFU measurements. Simulations were extended to 24 hours to capture the long-term steady-state behaviour of the bacterial populations beyond the experimentally measured growth window. **(B)** Identification of a sub-inhibitory "Stealth Window" (gray) where host pathology is resolved despite the persistence of a viable bacterial reservoir. The stealth window represents the parameter region where Σ exceeds baseline virulent infection levels. The shaded “Stealth Window” denotes the antibiotic concentration range where the Stealth Index (Σ) exceeds that of classical infection, indicating persistence of bacterial populations with reduced host pathology. **(C)** Predicted Stealth Index (Σ) demonstrating optimized bacterial load relative to host damage across an antibiotic gradient. At high antibiotic concentrations both populations decline; however, the proportional reduction in host pathology exceeds the reduction in total bacterial load, producing an increased Stealth Index. This reflects the conceptual nature of the metric rather than a strictly predictive quantity. **(D)** Summary of model parameters illustrating the sacrifice of acute virulence (α) for enhanced antibiotic survival (δ) in the L-form state. Parameter ranges were selected to qualitatively reproduce experimentally observed growth inhibition and timing of morphological transition rather than to achieve quantitative parameter estimation.

## Discussion

Shigellosis remains a major global health threat, particularly in low- and middle-income countries, and the global rise of extensively drug-resistant (XDR) strains resistant, to frontline antibiotics like cephalosporins is a pressing concern. Antibiotic resistance is a critical factor in bacterial survival, but beyond genetic adaptations, some bacteria employ morphological transformations, such as transitioning into cell wall-deficient L-forms, to endure antimicrobial pressure. In this context, we investigated the interplay between antibiotic resistance, L-form transition, and virulence in a clinical MDR *Shigella sonnei* isolate, HK8. Our findings compellingly demonstrate that while this pathogen is formidably equipped for acute infection, its primary survival strategy under β-lactam pressure—transitioning to an L-form—mandates a sacrifice of its classical virulence. This trade-off is characterized by a significant reduction in the virulence coefficient (α) in exchange for enhanced tolerance (δ) to cell-wall active agents. This elucidation of a critical virulence-survival trade-off has significant implications for understanding bacterial persistence and the challenges of treating MDR infections.

Our initial genomic and phenotypic analyses established HK8 as a clinically relevant and highly adapted pathogen. This strain exhibited substantial genetic similarity to *Escherichia coli*, consistent with their common evolutionary ancestor, a trait that may extend to shared mechanisms of resilience. The isolate was confirmed to be multidrug-resistant, with clear resistance to ceftriaxone. Our genomic investigation provided a direct molecular blueprint for this phenotype, identifying key point mutations in genes like *gyrA* and *PBP3*, alongside mutations in the multidrug efflux regulator *marR*. The *in-silico* profiling further revealed a vast arsenal of virulence factors, positioning HK8 as a canonical invasive pathogen. This comprehensive characterization underscores the dual threat posed by modern clinical isolates: high-level, genetically encoded resistance coupled with a full complement of virulence determinants.

A central finding was the bacterium’s multifaceted response to sublethal ceftriaxone exposure. The antibiotic induced a profound morphological shift into a filamentous state, a well-documented consequence of PBP3 inhibition. Significantly, these filamentous forms exhibited a hyper-adhesive phenotype on RAW 264.7 macrophages. This observation suggests a pivotal, antibiotic-driven switch in host-interaction strategy. This aligns powerfully with recent evidence demonstrating that host cells, particularly phagocytes, can serve as a protective niche for bacterial L-form generation and survival, effectively shielding them from both antibiotics and immune surveillance [11]. It is also important to note that antibiotic pressure is not the sole inducer; host-derived lysozymes within macrophages can also facilitate this transition. The antibiotic-induced adhesion we observed may therefore represent a critical initial step in establishing a surface-associated or intracellular reservoir.

The functional validation of cell wall deficiency in *S. sonnei* was confirmed through differential membrane permeability assays, which revealed a distinct “permeability-reversion” phenotype. While vegetative cells with intact peptidoglycan exclude Propidium Iodide (PI), the ceftriaxone-induced L-forms demonstrated significant PI uptake, providing direct experimental evidence of a compromised or absent cell wall barrier. Crucially, our observation that these variants could resynthesize their cell wall and restore vegetative morphology within one hour of removing antibiotic pressure proves they are viable, metabolically active persistant cells rather than terminal or degenerative populations. This rapid reversibility highlights a significant clinical risk: while L-forms remain attenuated and “stealthy” under antibiotic stress, they retain the full potential to revert to a virulent state once the therapeutic pressure is withdrawn, facilitating recurrent shigellosis.

Although the exact statistical probability of *Shigella sonnei* L-form occurrence in clinical strains remains difficult to quantify due to inherent diagnostic limitations, their role in chronic shigellosis is increasingly recognized. L-forms are notoriously fastidious and typically fail to grow under standard laboratory conditions, leading to their consistent underreporting in clinical datasets. Research indicates that under physiological stress, *Shigella* can undergo this phenotypic switch to evade both β-lactam antibiotics and host immune detection [11, 14]. This mechanism is particularly relevant in immunocompromised individuals, where the failure to achieve complete bacterial clearance often correlates with the presence of such persistent, non-replicating variants. Consequently, addressing the L-form transition is critical for improving clinical outcomes in recalcitrant cases where real-world data suggest standard therapies resolve symptoms without achieving true bacterial sterilization.

The most striking finding, however, was the profound and unequivocal attenuation of virulence observed in the L-form variant across multiple rigorous *in vivo* models. Despite being derived from a highly pathogenic strain, the L-form was completely unable to cause significant disease or histopathology. This loss of acute pathogenicity is likely multifactorial. The absence of a rigid cell wall would almost certainly compromise the structural integrity and function of essential, membrane-spanning virulence machinery, particularly the T3SS injectisome [12]. Furthermore, the likely loss or modification of surface antigens like LPS – which are the first molecules to interact with host cells and are under tight regulatory control – would fail to trigger the potent inflammatory cascade that defines the pathology of acute shigellosis. This was evident in the Sereny test, where L-forms failed to induce keratoconjunctivitis, and in the ileal loop model, where L-form inoculation resulted in significantly lower pathology scores, with reduced epithelial shedding, preserved barrier integrity, and minimal PMN cell infiltration.

The transition of *Shigella sonnei* into cell-wall-deficient L-forms represents a sophisticated survival strategy that allows the pathogen to persist within the host despite intensive antibiotic therapy and immune pressure. Although the exact prevalence of these forms in clinical isolates is currently underreported due to their fastidious nature and the limitations of standard laboratory diagnostics, their role in chronic and recurrent shigellosis is increasingly recognized. Research indicates that under physiological stress, Shigella can undergo phenotypic switching, resulting in L-forms that are resistant to beta-lactam antibiotics and less visible to the host’s innate immune system. This mechanism is highly relevant for immunocompromised individuals, where the failure to achieve complete bacterial clearance often points to the presence of such persistent, non-replicating variants. Consequently, addressing the L-form transition is critical for developing more effective therapeutic strategies and improving clinical outcomes in recalcitrant cases of Shigella infection [13, 14, 15].

This dichotomy between antibiotic survival and attenuated virulence firmly positions the L-form as a compelling model for a "stealth" or "persister" phenotype, which may mechanistically explain many chronic or recurrent infections [16]. It provides a potential mechanism for the long-term asymptomatic carriage of *S. sonnei* that has been previously documented. L-forms have been increasingly implicated in a range of intractable human diseases where bacteria persist silently, such as recurrent UTIs and even bloodstream infections where they evade standard detection methods [17]. This persistence is compounded by their ability to revert to the native, virulent cell-walled form once antibiotic pressure is removed, facilitating relapse. The ability of the *Shigella* L-form to survive antibiotic pressure while causing minimal acute inflammation provides a powerful model for how this pathogen could establish a cryptic reservoir. This may also suggest a parallel to other pathogens that utilize host macrophages as vehicles for dissemination through pathways like the gut-liver axis, allowing for systemic persistence without overt symptoms [18].

Our mathematical framework provides a quantitative basis for the clinical challenge of antibiotic treatment failure in shigellosis. By modelling the transition kinetics, we show that β-lactam antibiotics act as a double-edged sword: they effectively dismantle the cell-walled machinery required for acute invasion (the virulence side of the trade-off) but simultaneously trigger a phenotypic switch into a resilient, cell-wall-deficient state (the survival side). The high stealth index identified in our model explains a significant clinical paradox: the apparent resolution of acute symptoms following antibiotic therapy despite the persistence of a viable bacterial reservoir. These findings raise the possibility that certain antibiotic exposure regimes may inadvertently promote phenotypic switching into persistent L-form reservoirs, resolving acute symptoms while permitting long-term bacterial survival. From an evolutionary perspective, this trade-off allows *S. sonnei* to avoid host immune clearance by shedding its most inflammatory antigens (like peptidoglycan and LPS components), effectively adopting a cryptic lifestyle. These findings emphasize that true therapeutic success should not be measured merely by the reduction of pathology, but by the elimination of the stealth reservoir, highlighting the need for diagnostic and therapeutic strategies capable of detecting and eliminating cell wall–deficient bacterial reservoirs that may otherwise evade conventional antimicrobial therapy.

Our results provide direct experimental evidence for a stark virulence-survival trade-off in *S. sonnei*. The intense selective pressure to survive a cell wall-active antibiotic forces a phenotypic switch that simultaneously dismantles the complex machinery required for acute, invasive disease. This finding is of high clinical relevance, as it strongly implies that standard antibiotic regimens may inadvertently select for a population of bacteria that, while not causing immediate symptoms, are poised for long-term persistence and subsequent relapse [11]. These persisters represent a major diagnostic and therapeutic challenge, as they are often non-culturable by standard methods and are intrinsically resistant to many of our most common drugs [16, 17].

In conclusion, our work demonstrates that a clinical MDR isolate of *S. sonnei* can effectively evade β-lactam antibiotics by transitioning into a viable L-form state, but this survival comes at the price of its acute virulence. This trade-off between antibiotic survival and attenuated virulence positions the L-form state as a stealth persistant phenotype, in which bacterial populations can remain viable within host tissues while minimizing inflammatory detection. The very mechanism used to eradicate the pathogen provides the evolutionary pressure for its persistent survival. These findings highlight the need for therapeutic strategies that target not only actively growing bacteria but also phenotypically distinct, cell wall–deficient reservoirs that may underlie persistent and recurrent infection. Together, these findings suggest that phenotypic plasticity, rather than genetic resistance alone, may play a central role in shaping treatment outcomes in multidrug-resistant enteric pathogens.

## Materials and Method

### Bacterial Culture and Conditions

*Shigella sonnei* strain PD552A (HK8), was initially revived from glycerol stock using Tryptic Soy agar (TSA, BD Difco, USA) and sub-cultured in either Tryptic Soy broth (TSB, BD Difco, USA) broth or TSA plates at 37°C under aerobic conditions following the protocols published previously [19]. Similar culture conditions were maintained for *Vibrio cholerae* (N1961). To induce and maintain L-form, TSB was prepared as previously described [20] with slight modification. Briefly, TSB was supplemented with 0.8 M sucrose, 20 mM MgCl_2_, 1 mM CaCl_2_, and 10% Fetal Bovine Serum (cat# A5670701, Gibco). The maintenance buffer used during confocal microscopy to resuspend the bacteria contained 0.5 M sucrose, 20 mM MgCl_2_, 1 mM CaCl_2_, 10 mM HEPES (cat# 63732, SRL, pH 7.2). L-form was maintained under 5 μg/ml of ceftriaxone.

### Genomic Library preparation and Submission

High-quality genomic DNA from *Shigella sonnei* PD552A (HK8) was isolated utilizing the Qiagen Mini DNA Isolation Kit in accordance with the manufacturer’s protocol. The concentration and purity of the extracted DNA were evaluated spectrophotometrically using a NanoDrop system (Thermo Fisher Scientific), ensuring suitability for downstream applications. Whole-genome sequencing was performed on an Illumina platform, generating paired-end FASTQ reads.

Initial quality control and preprocessing of the raw reads were conducted using Fastp v0.23.4, enabling efficient filtering, adapter trimming, and removal of low-quality bases. The curated high-quality reads were then assembled *de novo* using Unicycler v0.4.4 under default parameters to generate draft genome assemblies. For improved contiguity and structural accuracy, reference-guided scaffolding was subsequently performed using RagTag v2.1.0, employing closely related *Shigella* reference genomes as templates. To facilitate accurate taxonomic classification, 16S rRNA gene fragments were extracted from the assembled contigs via the ContEST16S pipeline integrated within the EzBioCloud platform. Taxonomic affiliation was inferred based on sequence homology against the EzBioCloud 16S reference database. For comprehensive genome-based phylogenomic analysis and strain-level classification, the assembled genome was submitted to the Type Strain Genome Server (TYGS) (https://tygs.dsmz.de/). The resulting phylogenetic tree was rendered and annotated using Interactive Tree Of Life (iTOL), providing high-resolution insights into the evolutionary positioning of the isolate. Additionally, circular genome visualization was constructed using GenoVi v0.4.3 to represent key genomic features and architecture. The results have been submitted to NCBI under the accession number JBISBP000000000.

### Antimicrobial Susceptibility Test

Antimicrobial susceptibility was determined using the Kirby-Bauer disk diffusion method on Mueller-Hinton agar following Clinical and Laboratory Standards Institute (CLSI) guidelines [21]. Discs containing Chloramphenicol (30 μg), Ampicillin (10 μg), Kanamycin (30 μg), Neomycin (30 μg), Ciprofloxacin (5 μg), Furazolidone (100 μg), Ceftriaxone (30 μg), Erythromycin (15 μg), Sulphathiazole (300 μg), Levofloxacin (5 μg), Doxycycline (30 μg), Co-Trimoxazole (25 μg), Tetracycline (30 μg), Azithromycin (15 μg) were used. Zone diameters were measured after 18 hours of incubation at 37°C.

### Minimum Inhibitory Concentration (MIC) Determination

MIC values were assessed by the broth micro-dilution method according to CLSI protocols [22]. Serial two-fold dilutions of antibiotics were prepared in Mueller-Hinton broth (BD, Difco, USA) in 96-well microtiter plates. A gradient concentration of ceftriaxone starting from 10_μ_g to 60_μ_g was evaluated for MIC. The lowest concentration that completely inhibited visible bacterial growth after 18 hours at 37°C was recorded as the MIC.

### Growth Kinetics

Bacterial growth kinetics was analyzed by monitoring OD600 at 1-hour intervals for eight hours. Cultures were incubated in a shaking incubator (200 rpm) at 37°C. Growth curves were plotted, and doubling times were calculated during the exponential growth phase [23].

### Confocal Microscopy

#### L-form Characterization

For characterization, four groups of bacteria were maintained, namely, wild type (WT), L-form, revert form, and 70% ethanol treated. Initially they were grown in TSB for four hours without any supplements. A secondary culture was started after four hours at 1:100 dilution in the TSB supplemented media. Thirty minutes after starting the secondary culture, the L-form inducing group and revert group were supplemented with 5 μg/ml final concentration of ceftriaxone for four hours. Ceftriaxone was then removed from the revert group for an hour. Bacteria were then resuspended in the maintenance buffer mentioned above for further processing. Component A and component B from LIVE/DEAD BacLight Bacterial Viability (Invitrogen, cat# L7012) kit was mixed in 1:1 ratio and 3 μl of this mixture was added to each group of bacteria and kept for 15 minutes in room temperature. Bacteria were then washed and trapped between a glass slide and 18 mm^2^ glass cover slip for visualization under Zeiss LSM confocal microscope.

### Adhesion Assay

For adhesion assay, RAW 264.7 cells were grown on glass coverslips and infected. Post-infection, cells were fixed with 4% paraformaldehyde, permeabilized with 0.1% Triton X-100, and stained with in-house developed polyclonal anti-*Shigella* antibody and fluorescent secondary antibody (AlexaFluor488, Abcam). Samples were imaged using a Zeiss LSM confocal microscope [24].

### Preparation of Bacterial Inocula for Animal Studies

For all *in vivo* experiments, the inocula for the wild-type and L-form groups were standardized to ensure equivalent bacterial counts (1 x 10^7^ or 1 x 10^8^ CFU as specified), with concentrations verified by optical density (OD600) and serial dilution plating prior to administration.

### Animal ethics statement

All experimental protocols were performed in strict adherence to the guidelines of the Committee for the Control and Supervision of Experiments on Animals (CCSEA), having received prior approval from the Institutional Animal Ethics Committee (IAEC Approval No. Apo/80/06/05/2011, PRO/183/- Jan 2021-24, PRO/184/ - Jan 2021-24 and PRO/2034/Dec/2025 – 2028). The animals were maintained under a 12-hour light/dark cycle, a constant temperature of 22 ± 2°C, with standard chow and water *ad libitum*.

### Sereny Test in Guinea Pig

To assess *in vivo* virulence, the Sereny test was performed by instilling 1 x 10^7^ CFU of respective bacteria (either WT or L-form) into the conjunctival sac of guinea pigs (n = 3 per group). Animals were monitored daily for 5 days for signs of keratoconjunctivitis. Inflammation and mucopurulent discharge were recorded as positive responses. All procedures were conducted under approved ethical protocols [25].

### Intraperitoneal (IP) Infection in BALB/c Mice

Female BALB/c mice (6–8 weeks old, n = 6 per group) were injected intraperitoneally with 1 x 10^8^ CFU of respective bacteria (either WT or L-form) in PBS. Control groups received sterile PBS. Mice were monitored for signs of morbidity, mortality, and body weight loss [26].

### Ileal Loop Assay and Histopathological Observations

The ileal loop assay was performed in guinea pigs (n = 3 per group) under anaesthesia as described previously [27]. Approximately 5 cm ileal segments were ligated and injected with 1000 μL of bacterial suspension (1 x 10^8^ CFU/mL). After 18 hours, the animals were euthanized. Tissue samples were fixed in 10% formalin, embedded in paraffin, sectioned, and stained with hematoxylin and eosin (H&E) for histopathological examination under light microscopy.

### Mathematical Modeling of Population Dynamics and Phenotypic Switching

To integrate our experimental observations into a quantitative framework, we developed a conceptual compartmental model describing the population dynamics and phenotypic switching of *Shigella sonnei* HK8 under β-lactam antibiotic pressure. The model captures the experimentally observed trade-off between acute virulence and antibiotic survival mediated through L-form transition.

Model Structure and Population Dynamics:

The model considers two bacterial states: the classical cell-walled form (B_c_) and the cell wall–deficient L-form (B_l_). Population dynamics were described using a system of ordinary differential equations (ODEs):

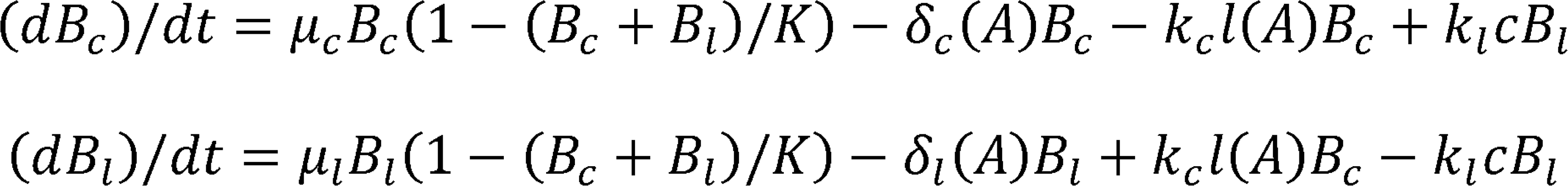

The logistic growth term reflects competition for a shared host-imposed carrying capacity K, which represents environmental limitations such as nutrient availability and immune pressure.

Antibiotic dependent processes:

Antibiotic concentration is represented by A.

The antibiotic-induced death rates were modeled as monotonic functions of antibiotic concentration:

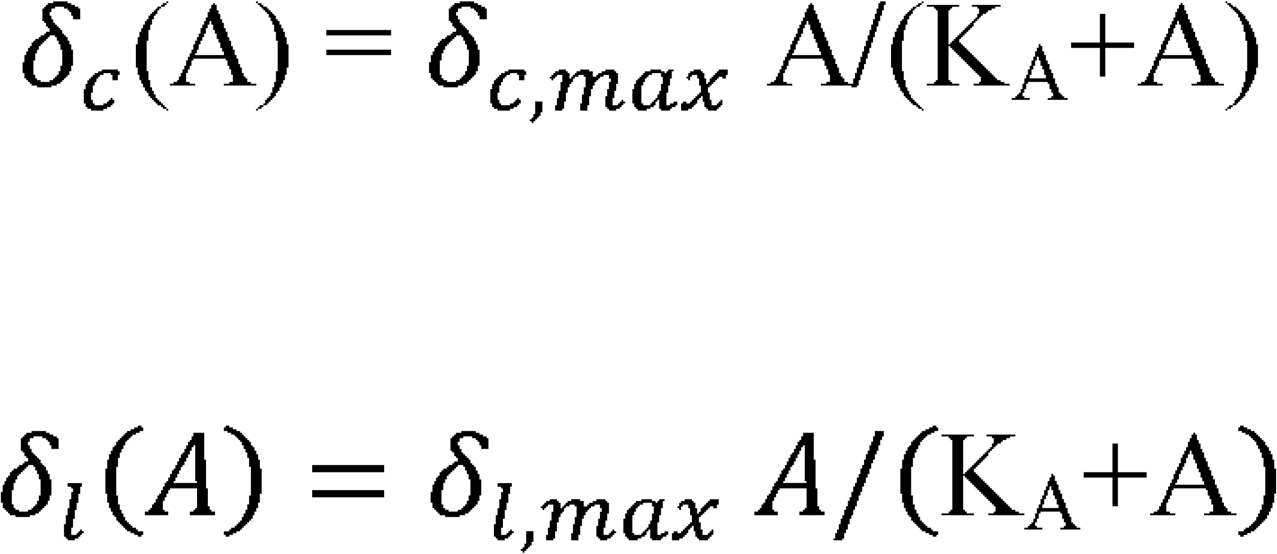

Where δ_c,max_ ≫ δ_l,max_, reflecting the well-established resistance of L-forms to cell wall-active antibiotics.

Phenotypic switching from classical cells to L-forms was modeled using a Hill-type function:

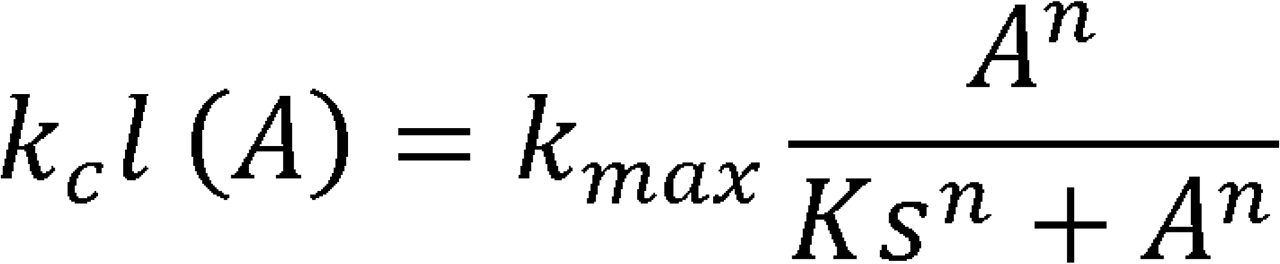

which captures the experimentally observed increase in filamentation and L-form transition with increasing ceftriaxone exposure.

The reverse transition rate k_lc_ represents spontaneous reversion of L-forms to the classical state upon removal of antibiotic stress, consistent with previous reports describing L-form reversibility.

Parameter Definitions

| Parameter | Description |
| --- | --- |
| $\mu_c$ | Growth rate of classical bacteria |
| $\mu_l$ | Growth rate of L-form bacteria |
| $K$ | Host carrying capacity |
| $\delta_c(A)$ | Antibiotic-dependent death rate of classical cells |
| $\delta_l(A)$ | Antibiotic-dependent death rate of L-forms |
| $k_{cl}(A)$ | Antibiotic-induced transition rate from classical cells to L-forms |
| $k_{lc}$ | Reversion rate from L-forms to classical cells |
| $A$ | Antibiotic concentration |

Model parameters were constrained using experimentally observed growth inhibition and time-resolved morphological transitions measured during ceftriaxone exposure (Fig. 6). Parameter ranges were constrained using experimentally observed growth curves and qualitative transition timing rather than formal parameter fitting, as the model is intended to provide a conceptual framework for interpreting the virulence–survival trade-off rather than a strictly predictive quantitative system. All simulations were performed across biologically plausible parameter ranges consistent with experimentally observed growth inhibition and phenotypic transition behavior.

Quantification of Host Pathology and the Stealth Phenotype:

To connect bacterial population dynamics with host disease severity, we defined a host pathology function:

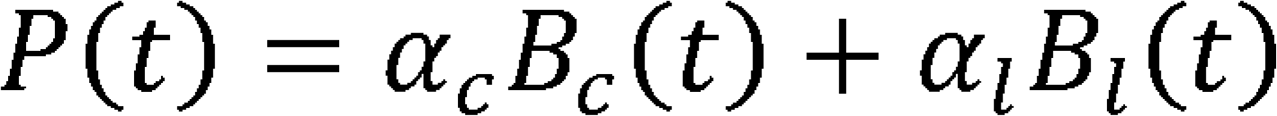

where

- α_c_ represents the virulence contribution of classical bacteria
- α_l_ represents the virulence contribution of L-forms

Consistent with our animal infection models (Figs. 4–5), classical bacteria exhibit substantially higher pathogenic potential: a_c_ ≫ a_l_

To quantify the persistence-virulence trade-off, we defined a Stealth Index, Σ *= (Bc + Bl)/P*. This formulation captures the relationship between pathogen burden and host damage, a commonly used conceptual framework in infection biology to distinguish between highly virulent pathogens that cause severe pathology and persistent organisms that maintain high population levels while eliciting limited host responses. This index represents the ratio of bacterial burden to host pathology.

Higher values of Σ correspond to conditions where bacterial populations persist with minimal host damage, a scenario consistent with L-form–dominated infections observed in our experimental models. Importantly, this index is used here as a conceptual metric rather than a strictly predictive mathematical quantity.

### Conceptual Implications

Model simulations reveal the existence of a therapeutic tipping point at intermediate antibiotic concentrations. At these levels:

- classical virulent populations collapse due to antibiotic sensitivity
- L-form populations expand through antibiotic-induced phenotypic switching
- host pathology is reduced due to the diminished virulence of L-forms

This dynamic explains how antibiotic therapy may resolve acute symptoms while simultaneously permitting persistence of a cryptic L-form reservoir capable of reversion and recurrent infection.

### Statistical Analysis

All experiments were performed in triplicate unless stated otherwise. Data are expressed as mean ± standard deviation (SD). Statistical analyses were conducted using GraphPad Prism v9. Graphs were prepared in GraphPad Prism v9 or R v4.5. Comparisons between groups were made using Student’s t-test or one-way ANOVA with Tukey’s post hoc test. A p-value < 0.05 was considered statistically significant.

## Acknowledgements

The authors thank Mr. Pritam Nandy, Ms. Samima Sultana, Mr. Suhasit Ranjan Ghosh, Mr. Subrata Singha, and Ms. Usha Hansdah for their technical assistance.

## Funding agency

The work was supported by the National Institute of Infectious Diseases, Japan in collaboration with ICMR-National Institute for Research in Bacterial Infections, Kolkata. DRH was funded by DBT-RRF India. UGC, India provided fellowship to Sanjib Das under CSIR-UGC-NFSC scheme [SANJIB DAS/3363/CSIR-UGC NET JUNE 2018] and Arindam Mukherjee under CSIR-UGC-NET Scheme [231620010720].

## Data availability

All data will be made available upon request.

## Author Contributions

**SD:** Conceptualization, design, investigation, data curation, methodology and writing; **AM:** Design, investigation, data curation, methodology, reviewing and editing of manuscript; **PH, SB, SM, NR, AD, SB, MM, UM:** Investigation and review; **JM, JHW, AKM, SD:** Review and editing; **DRH:** Design, investigation, data curation, writing, reviewing and editing of manuscript; **HK:** Conceptualization, design, funding acquisition.

## Notes

### Competing Interest Statement

The authors have declared no competing interest.

### Summary of Updates

Uploaded the current version of the manuscript.

