## Supplemental materials for "Third generation cephalosporin-induced L-form transition in *Shigella sonnei* reveals a virulence-survival trade-off underlying persistent infection"

**Supplementary Fig. S1.**

**A.**

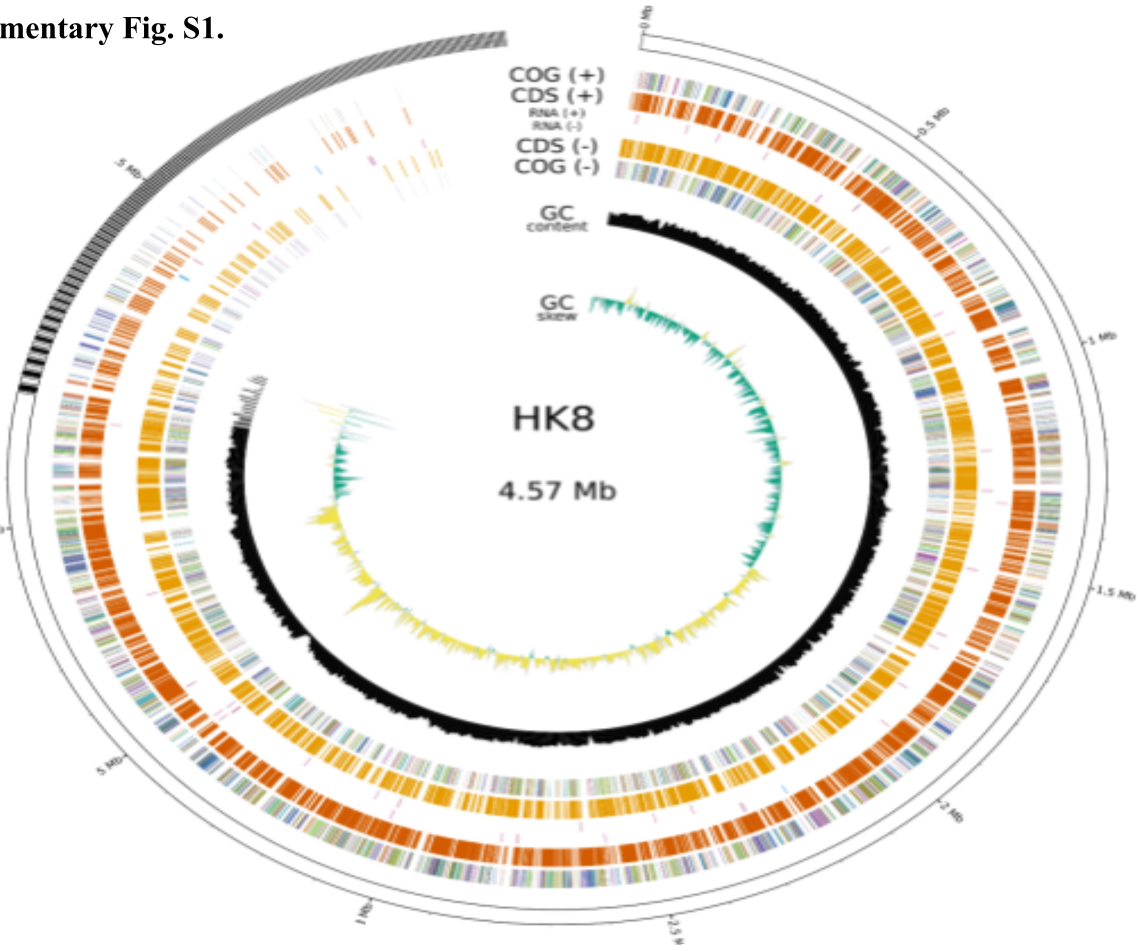


**B.**

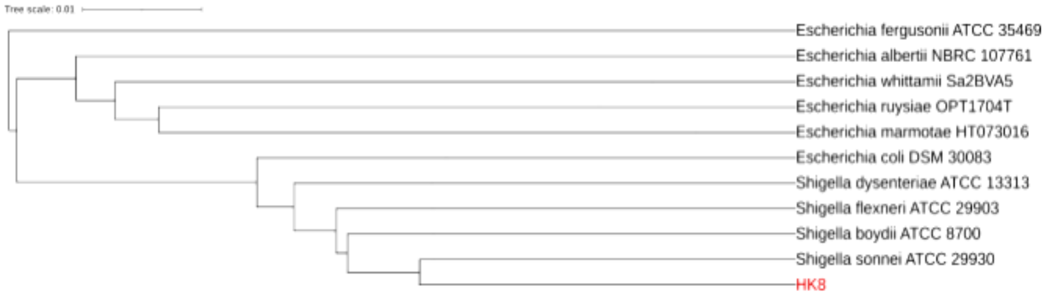


**Supplemental Fig. S1. Genomic Characterization, Phylogenetic Placement, and Virulence Profile of *Shigella sonnei* HK8.** **(A)** Circular map of the 4.57 Mb chromosome of *S. sonnei* HK8. The concentric rings, from the outside in, represent: (1-2) COG functional categories and coding sequences (CDS) on the positive strand; (3) RNA genes; (4-5) CDS and COG categories on the negative strand; (6) GC content relative to the mean; and (7) GC skew [(G-C)/(G+C)]. **(B)** Whole-genome phylogenomic tree illustrating the evolutionary position of *S. sonnei* HK8. The analysis, performed using the Type Strain Genome Server (TYGS), places HK8 (highlighted in red) directly adjacent to the *Shigella sonnei* ATCC 29930 type strain, confirming its species identity. The tree scale indicates the evolutionary distance.

### Third generation cephalosporin-induced L-form transition in *Shigella sonnei* reveals a virulence-survival trade-off underlying persistent infection

Sanjib Das[1], Arindam Mukherjee[1], Prolay Halder[1], Soumalya Banerjee[1], Supriya Mandal[1], Nivedita Roy[1], Ashis Debnath[1], Sreejani Banerjee[1], Manjistha Manna[1], Upama Mandal[1], Jiro Mitobe[2], Jeffery H. Withey[3], Asish Kumar Mukhopadhyay[1], Santasabuj Das[1], Debaki Ranjan Howlader[1]*, Hemanta Koley[1]*

[1]Division of Bacteriology, ICMR-National Institute for Research in Bacterial Infections, P-33, CIT Road, Scheme-XM, Beliaghata, Kolkata-700010, India

[2] Department of Bacteriology, National Institute of Infectious Diseases (NIID), Tokyo, Japan

[3] Department of Biochemistry, Microbiology and Immunology, Wayne State University School of Medicine, Detroit, Michigan, USA

**Keywords:** *Shigella sonnei*, Ceftriaxone, L-form bacteria, Animal models, chronic infection.

**Supplementary table ST1: Antibiotic profile of PD552A (HK8)**

| Antibiotic | Disc Potency (μg) | Observed Zone Diameter (mm) | Preliminary Interpretation |
|---|---|---|---|
| Doxycycline | 30 | 20±4 | Intermediate/Susceptible |
| Tetracycline | 30 | 22±3 | Susceptible |
| Chloramphenicol | 30 | 6±1 | Resistant |
| Ampicillin | 10 | 6±0 | Resistant |
| Erythromycin | 15 | 14±5 | Intermediate/Resistant |
| Azithromycin | 15 | 19±3 | Susceptible |
| Co-Trimoxazole | 25 | 18±2 | Intermediate/Susceptible |
| Sulphamethizole | 300 | 17±4 | Intermediate/Resistant |
| Kanamycin | 30 | 15±1 | Resistant |
| Neomycin | 30 | 15±2 | Resistant |
| Ciprofloxacin | 5 | 6±2 | Resistant |
| Furazolidone | 100 | 6±0 | Resistant |
| Levofloxacin | 5 | 12±1 | Intermediate/Resistant |
| Ceftriaxone | 30 | 13±1 | Resistant |

**Supplementary table ST1: Antibiotic profile of PD552A (HK8):** Antibiotic disc diffusion assay was performed for *Shigella sonnei* PD552A. The interpretations were made following CLSI guidelines (https://clsi.org/shop/standards/m100/).

**Supplementary table ST2: Effect of point mutations in respective genes and on antibiotic sensitivity**

| **ORF_ID** | **ARG location** | **SNPs** | **AMR subclass** | **Corresponding Genes** |
|---|---|---|---|---|
| 4416 | 234 | R>F | elfamycin antibiotic | *tuf* |
| 3035 | 448 | E>K | fosfomycin | *glpT* |
| 3045 | 87 | D>G | triclosan | *gyrA* |
| 3789 | 137, 103 | Y>H G>S | cephalosporin; fluoroquinolone antibiotic; glycylcycline; penam; phenicol antibiotic; rifamycin antibiotic; tetracycline antibiotic; triclosan | *marR* |
| 986 | 350, 357 | D>N S>N | carbapenem; cephalosporin; cephamycin; monobactam; penam | *PBP3* |

**Supplementary table 2: Effect of point mutations in respective genes and on antibiotic sensitivity:** Resistant genes and their phenotypic effects were shown indicating the role of point mutations in the induction of antimicrobial resistance.
